# CLASSIFICATION OF PROKARYOTIC DNA METHYLTRANSFERASES BY CATALYTIC DOMAIN TOPOLOGY

**DOI:** 10.1101/2023.12.13.571470

**Authors:** M. Samokhina, A. Alexeevski

## Abstract

Prokaryotic DNA methyltransferases (MTases) are essential enzymes that play a crucial role in restriction-modification systems, preventing the degradation of self DNA by restriction enzymes. In addition, they contribute to the regulation of gene expression, DNA repair, DNA replication, and other processes.

Here, we propose a novel classification of prokaryotic MTases based on the 3D topology of the catalytic domain, which addresses the limitations of the previous scheme developed by Malone and Bujnicki. The latter, which grouped MTases into categories such as α, β, γ, etc., has been in use for over 20 years.

We identified seven structural classes (A–G) of MTases by analyzing 38 experimentally determined 3D structures and several AlphaFold-predicted models, and validated the classification against all reviewed DNA MTases from Swiss-Prot. The catalytic domains of each class exhibit homology not only in structure but also in conserved motifs.

Our classification system provides a framework for understanding the diversity and evolution of prokaryotic DNA MTases, with applications in genomic annotation and evolutionary studies.

## INTRODUCTION

DNA methylation at the C5 position of cytosine was first discovered in bacteria in 1925 [1], marking nearly a century of research into genome methylation and the enzymes responsible for this modification. These enzymes, DNA methyltransferases (MTases), were initially identified as components of restriction-modification (R-M) systems [1] and have since been recognized for their roles in diverse biological processes, including other bacterial immune mechanisms, cell cycle regulation, gene expression, and DNA replication [2,3].

MTases are typically classified using three complementary approaches: functional classification, classification by association with restriction-modification systems, and sequence similarity classification.

**Functional classification** categorizes MTases according to the specific chemical reactions they catalyze and the resulting products. This approach defines three major enzyme classes [4]:

• **C5-MTases**: Produce C5-methylcytosine (m5C; EC 2.1.1.37)

• **N6-MTases**: Produce N6-methyladenine (m6A; EC 2.1.1.72)

• **N4-MTases**: Produce N4-methylcytosine (m4C; EC 2.1.1.113)

These three types of DNA methylation —m6A, m4C, and m5C— are now well-established [5,6]. MTases from all three classes are found in bacteria and archaea, with N6-MTases being the most prevalent [7]. In eukaryotes, C5-MTases that methylate cytosine in CG dinucleotides are predominant [8]. The quantitative distribution of these MTase types across domains of life, based on UniProt data, is presented in Table 1.

**Table 1.**
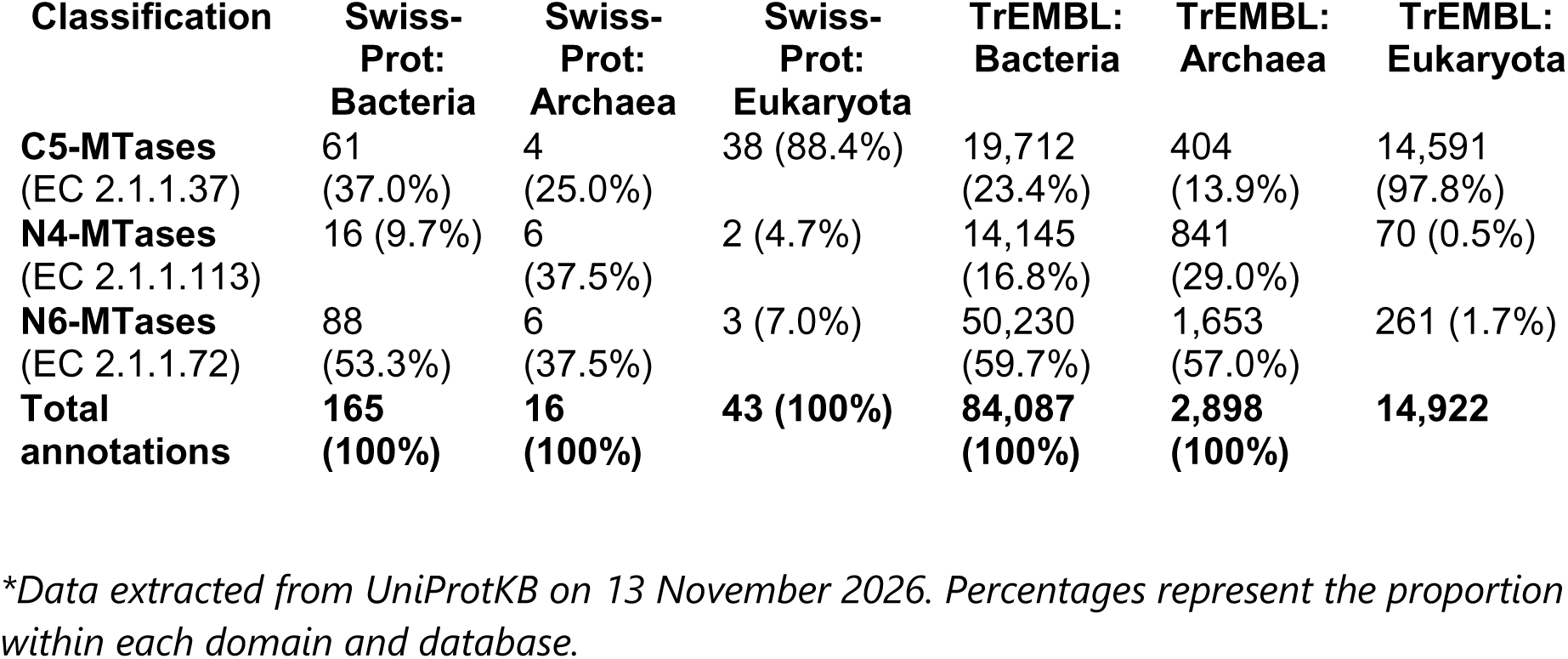
Distribution of DNA methyltransferases in UniProtKB*.

| Classification | Swiss-Prot: Bacteria | Swiss-Prot: Archaea | Swiss-Prot: Eukaryota | TrEMBL: Bacteria | TrEMBL: Archaea | TrEMBL: Eukaryota |
|---|---|---|---|---|---|---|
| **C5-MTases** (EC 2.1.1.37) | 61 (37.0%) | 4 (25.0%) | 38 (88.4%) | 19,712 (23.4%) | 404 (13.9%) | 14,591 (97.8%) |
| **N4-MTases** (EC 2.1.1.113) | 16 (9.7%) | 6 (37.5%) | 2 (4.7%) | 14,145 (16.8%) | 841 (29.0%) | 70 (0.5%) |
| **N6-MTases** (EC 2.1.1.72) | 88 (53.3%) | 6 (37.5%) | 3 (7.0%) | 50,230 (59.7%) | 1,653 (57.0%) | 261 (1.7%) |
| **Total annotations** | **165 (100%)** | **16 (100%)** | **43 (100%)** | **84,087 (100%)** | **2,898 (100%)** | **14,922** |

*\*Data extracted from UniProtKB on 13 November 2026. Percentages represent the proportion within each domain and database.*

**Classification by association with R-M systems** groups MTases based on their role within the restriction-modification machinery, a key prokaryotic defense system. MTases are integral components of Type I, II, and III R-M systems. In Type II systems, the MTase is typically a standalone enzyme, while in Type I and III systems, it acts as a subunit within a multi-enzyme complex [9]. A notable variant is Type IIG systems, where the genes for the restriction enzyme and the MTase are fused, resulting in a single polypeptide chain (RM proteins) [10]. Furthermore, a significant number of "orphan" MTases exist that are not associated with any restriction enzyme and are involved solely in regulatory functions, such as controlling gene expression and DNA replication [11].

**Sequence similarity classification** was introduced in 1995, when nine conserved motifs were identified in known MTases. Based on the order of these motifs and the position of the target recognition domain (TRD), MTases were categorized into four groups (α, β, γ, and a group for C5-MTases) [12]. This classification was later formalized into an evolutionary model involving circular permutations, which proposed six possible classes (α, β, γ, δ, ε, ζ) by accounting for different linear arrangements of the conserved subdomains. Among these, the α, β, γ, and ζ classes were confirmed in experimentally characterized MTases, while the δ and ε classes remained hypothetical [13].

Since 2002, the number of known MTase sequences has expanded dramatically, from fewer than 100 to over ten thousand, with this information aggregated in the Restriction Enzyme Database (REBASE) [14]. However, determining the precise number of proteins classified according to conserved motifs is challenging, because detection methods for all defined motifs and the TRD are limited. Of the nine motifs described, motif I (the SAM-binding motif) and motif IV (the catalytic motif) are the most reliably identified in REBASE entries; however, many MTase sequences lack any detectable canonical motifs due to high sequence divergence. Among the nine motifs described, motif I (the SAM-binding motif) and motif IV (the catalytic motif) can be found in a number REBASE entries, yet many MTases have no detected motifs at all.

In this work, we present a classification system for prokaryotic MTases based on the 3D topology of their catalytic domain, using a contemporary dataset from REBASE. We developed an algorithm that automatically assigns MTase structures to their topological class. We also provide a web application for visualizing classification results and, based on our findings, discuss the structural relationships between the identified topologies.

## MATERIALS AND METHODS

### Materials

#### MTase 3D Structures

We retrieved a list of 38 MTases with experimentally determined 3D structures from REBASE (as of July 31, 2026; see Table S1a). Notably, five of these MTases contain not only the methyltransferase catalytic domain but also the catalytic domains of restriction endonucleases. These dual-function proteins are designated with the prefix "RM" in REBASE.

For topologies not represented among experimentally determined structures, we used predicted models from the AlphaFold database (https://alphafold.ebi.ac.uk/).

#### Sequences from REBASE

We obtained MTase sequences from REBASE v408. We retained proteins that were included in the REBASE gold standard, were also present in UniProt, and had a predicted AlphaFold model in the AlphaFold database (see Table S1b).

To assess the coverage of the pipeline, it was additionally applied to all reviewed DNA MTases in Swiss-Prot (EC 2.1.1.37, 2.1.1.72, 2.1.1.113; Table S1e). Of the 181 proteins initially retrieved, 5 were excluded as gene fragments or non-MTase entries (Table S4), leaving 176 proteins for analysis.

## Methods

### Topology classification of Mtases

Secondary structure was assigned using DSSP (version 4.2.2). Beta-strands (DSSP code ’E’) were merged into continuous segments with a minimum length of three residues. Alpha-helices (codes ’H’, ’G’, ’I’) were merged with a minimum length of eight residues.

Catalytic motifs were searched within strand sequences using regular expressions. Default patterns were ‘[SND]P[PL][YFW]’ (covering NPPY, DPPY, SPPY, NPPF, DPPF, SPPF, DPPW, etc.) and ‘P[CS]’ (covering PC, PS). The strand containing a motif was designated S4 (S = strand).

For each structure, all strands belonging to the same chain as S4 were retained. A contact graph was built between strands based on the minimum atomic distance: two strands were considered adjacent if the distance between any of their atoms was below a threshold (CONTACT_DIST = 5.2 Å). Adjacent strands were then separated into N-terminal and C-terminal neighbours according to their position in the sequence relative to S4. The resulting per-structure strand assignments are provided in Table S1c (experimentally determined structures) and Table S1d (AlphaFold-predicted models).

### Structural comparison

Pairwise structural alignments were performed using US-align (version 20260920) with circular permutation mapping (-mm 3) and all-chain mode (-ter 0). For each pair of representative structures, the TM-score was extracted and assembled into a symmetric matrix (the maximum of the two directional TM-scores was used for each pair). Pairwise distances were calculated as 1 − TM-score, and hierarchical clustering was performed using UPGMA (average linkage, scipy.cluster.hierarchy). The resulting dendrogram was visualized with a distance threshold of 0.38, which separates the topologies into three superclusters. Structural superpositions for figures were generated using the jCE-CP algorithm (RCSB PDB Protein Comparison Tool, https://rcsb.org/alignment).

### Implementation

The automated classification pipeline was implemented in Python, and the interactive web application was developed using the Streamlit framework.

## RESULTS

The catalytic domains of MTases are thought to feature a conserved seven-stranded β-sheet (topology ↓↑↓↓↓↓↓) sandwiched between layers of α-helices [15]. To evaluate whether the entire spectrum of currently resolved MTase structures shares this architectural core, we implemented a geometry-driven automated search algorithm to computationally isolate the catalytic domain. Rather than relying on arbitrary manual truncation, our algorithm scans for catalytic motif IV and, once found, searches for the β-sheet around it.

To enable a standardized comparative analysis of these verified DNA MTase structures, we established a unified nomenclature based on a canonical spatial orientation (Figure 1a). In this standard view, the catalytic domain is oriented such that the catalytic core — a seven-stranded β-sheet — runs away from the viewer, with six parallel strands (blue triangles) and one antiparallel strand (inverted purple triangle). Within this framework, the α-helices located above the β-sheet are designated Hu1–Hu3, and those beneath it Hd1–Hd3 (where present). The core β-strands are numbered S1–S7 by their spatial position in the sheet, from left to right (Figure 1a); this numbering does not follow the order of strand appearance along the polypeptide chain (N→C). Figures 1b–1d show an example of the topology, 3D structure, and linear secondary-structure arrangement of M.CbeI.

**Figure 1.**
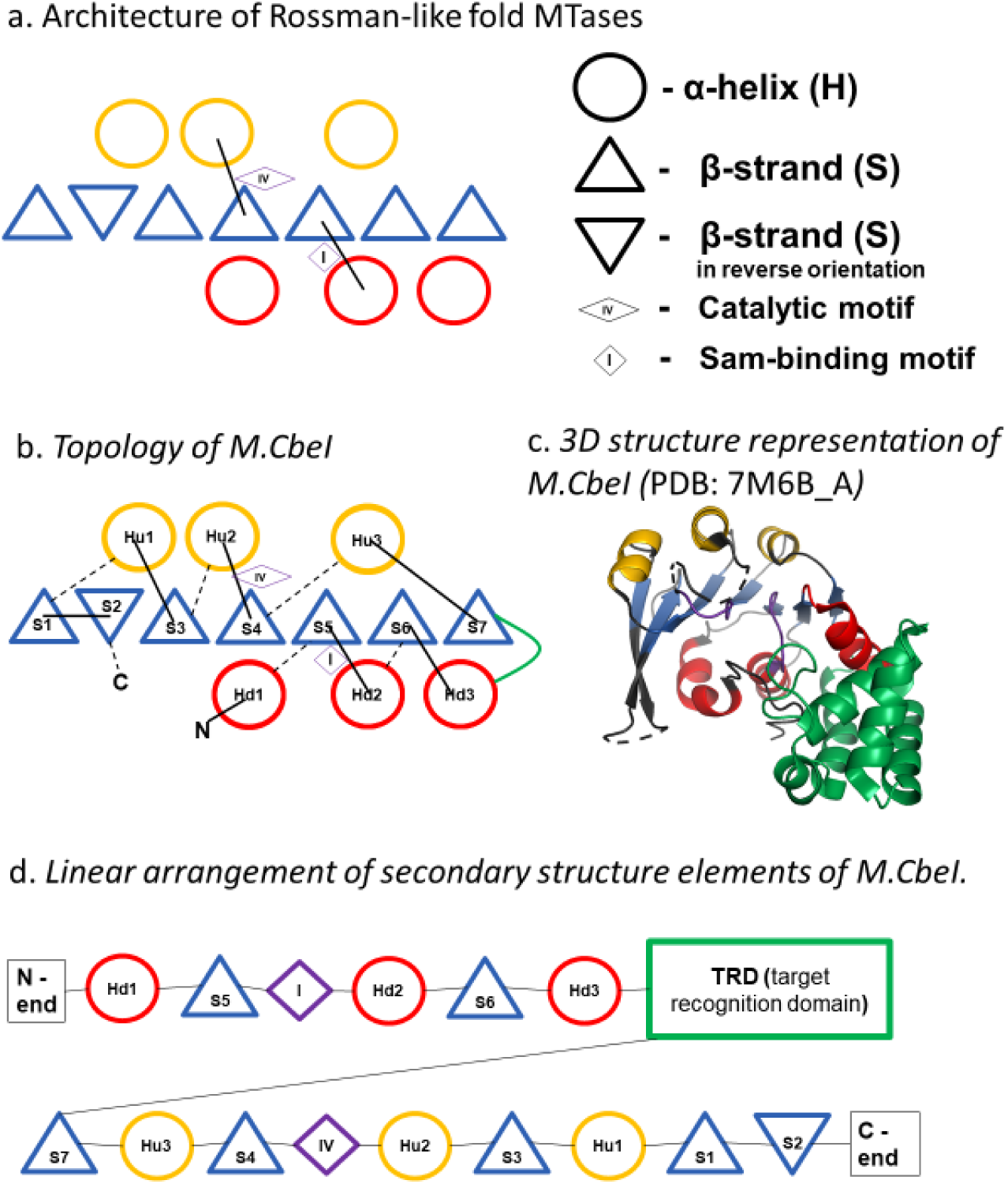
Topological scheme of the MTase catalytic domain. (a) Architecture of the Rossmann-like fold in MTases. The catalytic core is a seven-stranded β-sheet (triangles) sandwiched between layers of α-helices (circles). Blue triangles denote β-strands in the standard orientation; inverted blue triangles denote β-strands in the reverse orientation. Orange circles denote α-helices above the sheet; red circles denote α-helices below the sheet. The catalytic motif IV is marked with a diamond; the SAM-binding motif I is marked with a small circle. (b) Topology of M.CbeI (PDB 7M6B, chain A). The β-strands are numbered S1–S7 by their spatial position in the sheet, from left to right. Helices above the sheet are designated Hu1–Hu3; helices below the sheet are designated Hd1–Hd3. The N- and C-termini of the polypeptide chain are indicated. (c) 3D structure of the M.CbeI catalytic domain, colored according to the topology in (b). (d) Linear arrangement of the secondary structure elements of M.CbeI along the polypeptide chain (N→C), showing the order of β**-**strands, α-helices, the catalytic motif IV, and the target recognition domain (TRD).

### Topological classification of prokaryotic MTases based on 3D structures

As of July 31, 2026, 38 prokaryotic DNA MTases with resolved spatial structures are known, all of which share a catalytic core formed by a β-sheet sandwiched between layers of α-helices (see Table S2). Among them, five are RM proteins, meaning they also contain an endonuclease domain. Based on the topology of their catalytic domains — specifically, the order of β-strands, the presence of additional strands S0 and S-1, the gap between S6 and S7, and the arrangement of N-terminal and C-terminal helices — the 38 known MTase structures form four experimentally defined groups, designated as Topologies A, B, C, and D (Figure 2a). In addition, analysis of AlphaFold-predicted models revealed three further topologies (E, F, G) in proteins with experimentally confirmed MTase activity but no resolved structures (Figure 2b).

**Figure 2.**
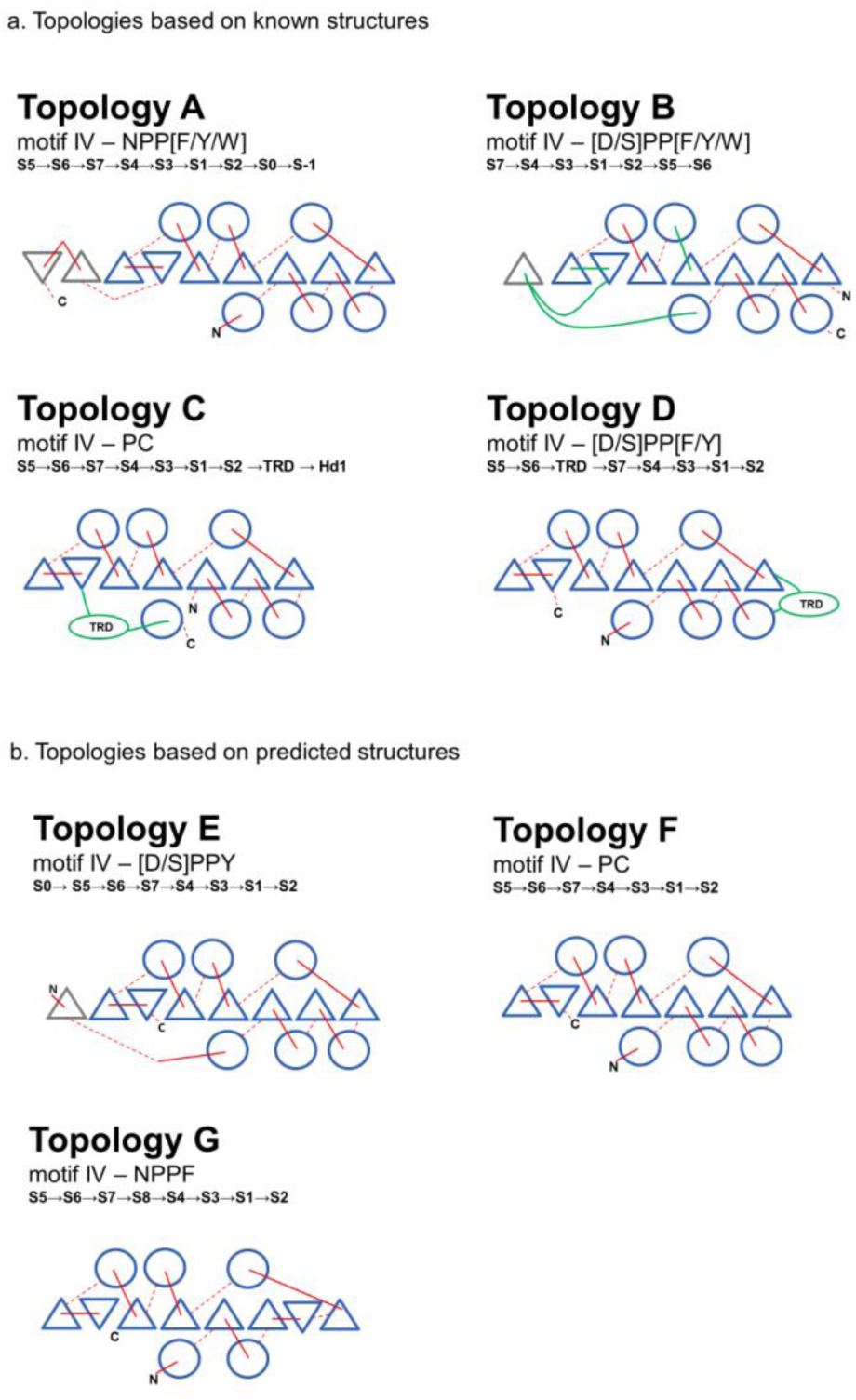
Topological schemes of the seven MTase classes identified in this study. (a) Topologies based on experimentally determined structures (A–D). (b) Topologies based on AlphaFold-predicted models (E–G). For each topology, the catalytic motif IV and the strand order are indicated above the scheme. Blue triangles denote β-strands in the standard orientation; inverted blue triangles denote β-strands in the reverse orientation. Blue circles denote α-helices. Solid red lines connect sequentially adjacent elements along the polypeptide chain; dashed red lines connect spatially adjacent elements. The N- and C-termini are marked. The target recognition domain (TRD) is shown in green. The strand order string for each topology is given below the name (e.g., S5→S6→S7→S4→S3→S1→S2→S0→S-1 for Topology A).

**Topology A** (motif IV: NPP[F/Y/W]; strand order: S5→S6→S7→S4→S3→S1→S2→S0→S-1) is the most abundant architecture among DNA MTases, comprising 18 of the 38 experimentally determined structures (Table S2) and 377 AlphaFold-predicted models (Table S1d). The catalytic domain of these proteins begins with the Hd1 element, and the core ends with S2. A defining feature of this topology is the presence of an S0 strand at the C-terminus, and most members also contain an additional S-1 strand. When S-1 is absent (e.g., M.EcoR124I, PDB 7BTP), the variant is designated A-truncated; when additional C-terminal strands (S-2, S-3) are present (e.g., M.BseCI, PDB 7QW8), it is designated A-extra. Notably, the A-truncated variant reflects incomplete resolution of the C-terminal region rather than a genuine deletion: for M.EcoR124I, the AlphaFold-predicted model of the same protein (UniProt P10484) contains the full set of strands, including S-1. Among the AlphaFold-predicted models assigned to topology A, 16 were classified manually. These models carried a motif typical of class A and did not exhibit the strand permutations or rearrangements characteristic of classes B, C, or D; however, they were not assigned by the automated pipeline and were therefore added to class A by exclusion. The catalytic motif in this topology is typically NPPF, NPPY, or NPPW. All five RM proteins in our dataset (RM.BpuSI, RM.DrdV, RM.LlaGI, RM.LlaBIII, RM.MmeI) belong exclusively to class A. Topology A MTases lack a TRD within the catalytic domain; instead, the target site is recognised by another subunit of the active complex (the S subunit),which is typical of Type I R-M systems and is sometimes C-terminally joined to the MTase. The size of the catalytic domain varies from 213 aa (M.TaqI) to 292 aa (RM.MmeI), with the exception of M.Sen23580I (361 aa), whose structure features extended insertions (loops of 35 and 42 residues) between secondary-structure elements; these insertions do not disrupt the overall domain topology.

**Topology B** (motif IV: [D/S]PP[F/Y/W]; strand order: S7→S4→S3→S1→S2→S0→S5→S6) is a highly abundant architecture among amino DNA MTases, comprising 10 of the 38 experimentally determined structures (Table S2) and a large set of AlphaFold-predicted models (Table S1d). This topology exhibits a permuted strand order: the sheet begins with strand S7 — sometimes preceded by additional N-terminal strands such as S8 or S9 — and terminates with S5 and S6 as the last two strands (in that order). Additional strands (S8, S9, S10) may be present at either terminus, and most members contain a strand S0 located within the target-recognition region (present in 162 of 166 AlphaFold models and in most experimentally determined structures; Tables S1d and S2); the presence of these extra strands does not affect the core permuted order. The catalytic motif in this class is most frequently DPPY, but variants such as DPPF, DPPW, SPPF, and SPPY also occur; the motif itself is not required for topological assignment. Topology B is characterized by the absence of the C-terminal S-1 strand that defines the canonical topology A.

Target interaction in topology B involves distinct recognition and binding elements (Figure 3). In M.EcoP15I (4ZCF) [16], an inserted TRD is located between strands S2 and S0. In contrast, M.CcrMI (6PBD) [17] and M.BceJIV (9C3S) [18] use structurally equivalent target recognition loops positioned at the same topological locations to engage the primary target site. However, their auxiliary DNA-binding elements differ: while nonspecific DNA binding in M.CcrMI is provided by a separate C-terminal DNA-binding domain, M.BceJIV instead possesses an outward-protruding domain composed of two short α-helices. Although previously annotated as a target recognition domain (TRD), this element in M.BceJIV does not drive sequence recognition; instead, it functions as a DNA-binding anchor that contributes to duplex capture and stabilization.

**Figure 3.**
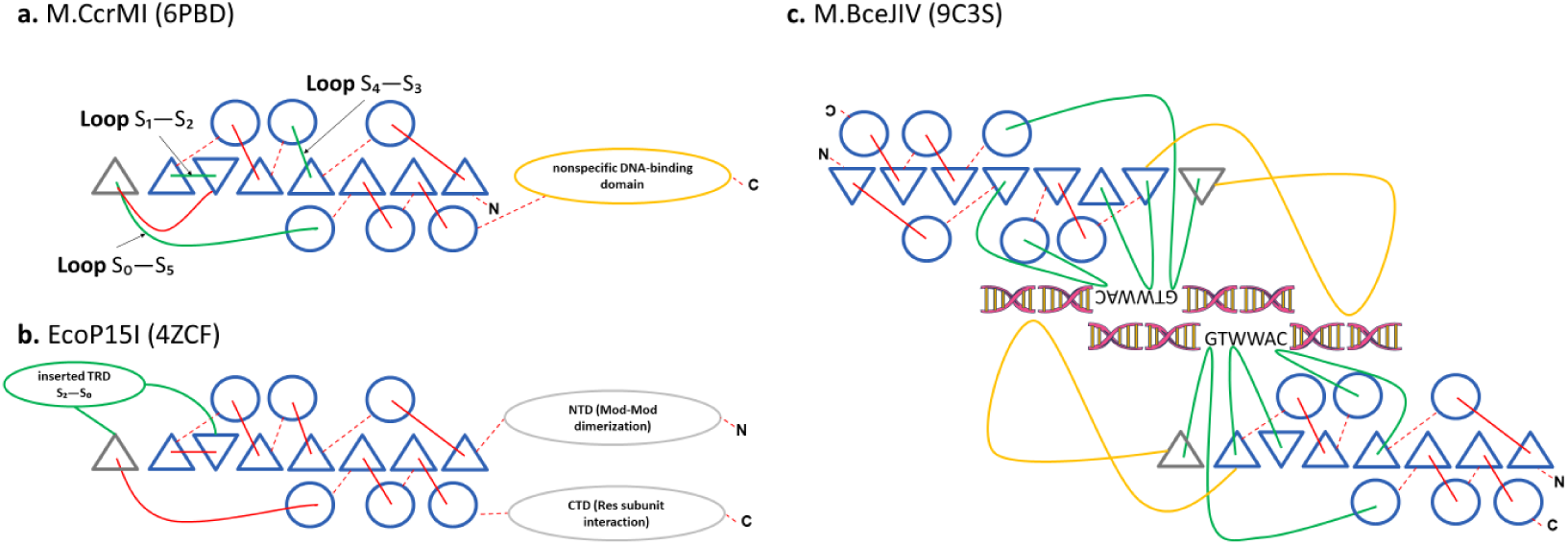
Target recognition and auxiliary DNA-binding elements in topology B. (a) Topological scheme of M.CcrMI (PDB 6PBD). The catalytic core is shown as in Figure 2; the target-recognition loops S1–S2, S4–S3, and S0–S5 are labelled. Nonspecific DNA binding is provided by a separate C-terminal DNA-binding domain (yellow oval). (b) Topological scheme of M.EcoP15I (PDB 4ZCF). An inserted target recognition domain (TRD) is located between strands S2 and S0 (green oval). The N-terminal domain (NTD) mediates Mod–Mod dimerization, and the C-terminal domain (CTD) mediates interaction with the Res subunit (grey ovals). (c) 3D structure of the M.BceJIV–DNA complex (PDB 9C3S). Two DNA duplexes are shown in yellow and pink. The outward-protruding domain, composed of two short α-helices, is highlighted. Green and yellow lines trace the polypeptide chain. Unlike the inserted TRD of M.EcoP15I, this element in M.BceJIV does not drive sequence recognition and instead functions as a DNA-binding anchor.

**Topology C** (motif IV: PC; strand order: S5→S6→S7→S4→S3→S1→S2) comprises five experimentally determined structures and is unique among experimentally determined structures exclusively associated with m5C methylation. This topology is characterized by the presence of a C-terminal helix and the absence of an N-terminal helix — a reversal of the helical arrangement seen in topologies A and B. The β-sheet is often reduced: several representatives lack strand S7 entirely (e.g., M.HhaI/9MHT, M.MpeI/8C57), and this reduction is recapitulated in 34 of the 150 AlphaFold-predicted models assigned to this topology (Table S1d). Topology C is further distinguished by the absence of the C-terminal S-1 strand in its canonical form, although 15 predicted models carry additional C-terminal strands (S0, S-1, S-2) and are designated C-extra. Unlike topologies A and B, the catalytic domains in topology C are the most extended, ranging from 312 aa (M.HhaI) to 441 aa (M.SflTDcmP); this size variation is primarily due to the C-terminal TRD, which in topology C occupies a strictly conserved position between structural elements S2 and Hd1.

**Topology D** (motif IV: [D/S]PP[F/Y]; strand order: S5→S6→S7→S4→S3→S1→S2) is represented by four experimentally determined structures — M.EcoKDam (2G1P), M1.DpnII (2DPM), M.EcoT4Dam (1YFJ), and M.CbeI (7M6B) — and a large set of AlphaFold-predicted models (Table S1d). In terms of secondary-structure organization, topology D resembles topology A, but differs by the absence of the C-terminal β-strands S0 and S-1 in its canonical form. A defining feature of this class is a large insertion: a compact TRD (60–80 aa) that is integrated between helix Hd3 and strand S7, ensuring structural compactness while maintaining functional activity. A small subset of predicted models carries additional C-terminal strands (S0, S-1) and is designated D-extra. Topology D is further characterized by the presence of an N-terminal helix and the absence of a C-terminal helix — the opposite of the helical arrangement seen in topology C. The catalytic domain size ranges from 231 aa (M.EcoT4Dam) to 255 aa (M.CbeI) among experimental structures. All four experimentally determined members of topology D are single-domain proteins. M.EcoKDam, M.EcoT4Dam, and M1.DpnII mediate m6A methylation, whereas M.CbeI is specific for m4C modification.

The catalytic motif is most frequently DPPY, but variants such as SPPY, SPPF, and DPPF also occur; as in other topologies, the motif itself is not required for topological assignment.

**In addition to the four experimentally defined topologies (A–D),** our analysis of AlphaFold-predicted models identified three new topologies (E, F, G) in proteins with experimentally confirmed MTase activity but no resolved structures.

**Topology E** (motif IV: [D/S]PPY; strand order: S0→S5→S6→S7→S4→S3→S1→S2) is represented by two AlphaFold-predicted models (Q8RNV6, Q5DX25) and includes the MTases M1.BcnI[19] and M4.BstF5I[20]. Topology E is structurally related to topology D — it is detected by the same set of B- and D-associated profiles and shares the S5→S6→S7→S4 →S3→S1→S2 core — but lacks the TRD insertion between Hd3 and S7. Consequently, the gap between strands S6 and S7 is much smaller (22–26 aa) than in topology D (≥100 aa). A defining feature of topology E is the presence of an N-terminal S0 strand. The catalytic motif in this class is DPPY or SPPY.

**Topology F** (motif IV: PC; strand order: S5→S6→S7→S4→S3→S1→S2) is represented by two AlphaFold-predicted models and includes two functionally characterized MTases: M2.Eco31I (Q8RNY6) and M.Esp3I (Q8RNY3) [21]. Topology F is related to topology C — both share the PC catalytic motif — but differs by the presence of an N-terminal helix and the absence of a C-terminal helix. Unlike some members of topology C, topology F contains a complete β-sheet (S5, S6, and S7 are all present). In one predicted model (M.Esp3I), an additional S8 strand is inserted between S6 and S7, but this does not affect the core fold. Notably, M.Esp3I contains two distinct MTase domains in a single polypeptide: an N-terminal domain with topology A (NPPY) and a C-terminal domain with topology F (PC).

**Topology G** (motif IV: NPPF; strand order: S5→S6→S7→S8→S4→S3→S1→S2) is represented by five AlphaFold-predicted models and includes M.EcoRI (P00472)[22–23] and M.MpnI (Q50290)[24], for which methyltransferase activity has been experimentally confirmed. In this topology, the segment between strands S6 and S7 adopts a β-strand conformation rather than an α-helix, and an additional strand S8 is present between S7 and S4. S7 is antiparallel to S6, while S8 is parallel to S4. Several features of the predicted models argue against this arrangement being a prediction artifact: the pattern is consistent across all five G models; the S6–S7 segment in M.MpnI is only 7 residues long, which is physically insufficient for an α-helix; the segment contains a proline that precludes a helical conformation; a hydrophobic core (Ile69, Phe71, Leu74, Tyr78) stabilizes the alternative fold; and a loop of 5–27 residues separates S7 and S8, indicating that they are two distinct strands rather than a single misannotated element. No experimental structure is currently available for any topology G member.

### Validation of the classification on reviewed MTases

To assess the coverage of the classification, we applied the pipeline to all reviewed DNA MTases in Swiss-Prot (EC 2.1.1.37, 2.1.1.72, 2.1.1.113). Of the 181 (Table S1e) proteins initially retrieved, 5 were excluded as gene fragments or non-MTase entries (Table S4), leaving 176 proteins for analysis. Of these, 168 were assigned to one of the seven topological classes, while 8 were not classified (Table S3). Three of the unclassified proteins carry non-canonical motif IV variants (DPQY, DIPY, NPHF) that fall outside the two motif families used by the parser, and five carry canonical motifs but were not classified because geometric constraints prevented assembly of the β-sheet around the motif. For all eight, the topological class was assigned manually, based on motif comparison and structural alignment with representative members of each class.

Notably, none of the reviewed Swiss-Prot MTases were assigned to topologies E or F. Both classes are represented exclusively by proteins that are absent from Swiss-Prot and were retrieved from REBASE and TrEMBL, and their assignment therefore relies entirely on AlphaFold-predicted structures.

### Structural relationships between topologies

To examine the structural relationships between the seven topologies, we performed pairwise structural alignments of representative members using jCE-CP, which accounts for circular permutations. Clustering based on TM-scores (1 − TM-score distance, UPGMA) revealed three major superclusters: (i) A + G, (ii) B + E, and (iii) C + F, with topology D forming a separate branch (Figure 4).

**Figure 4.**
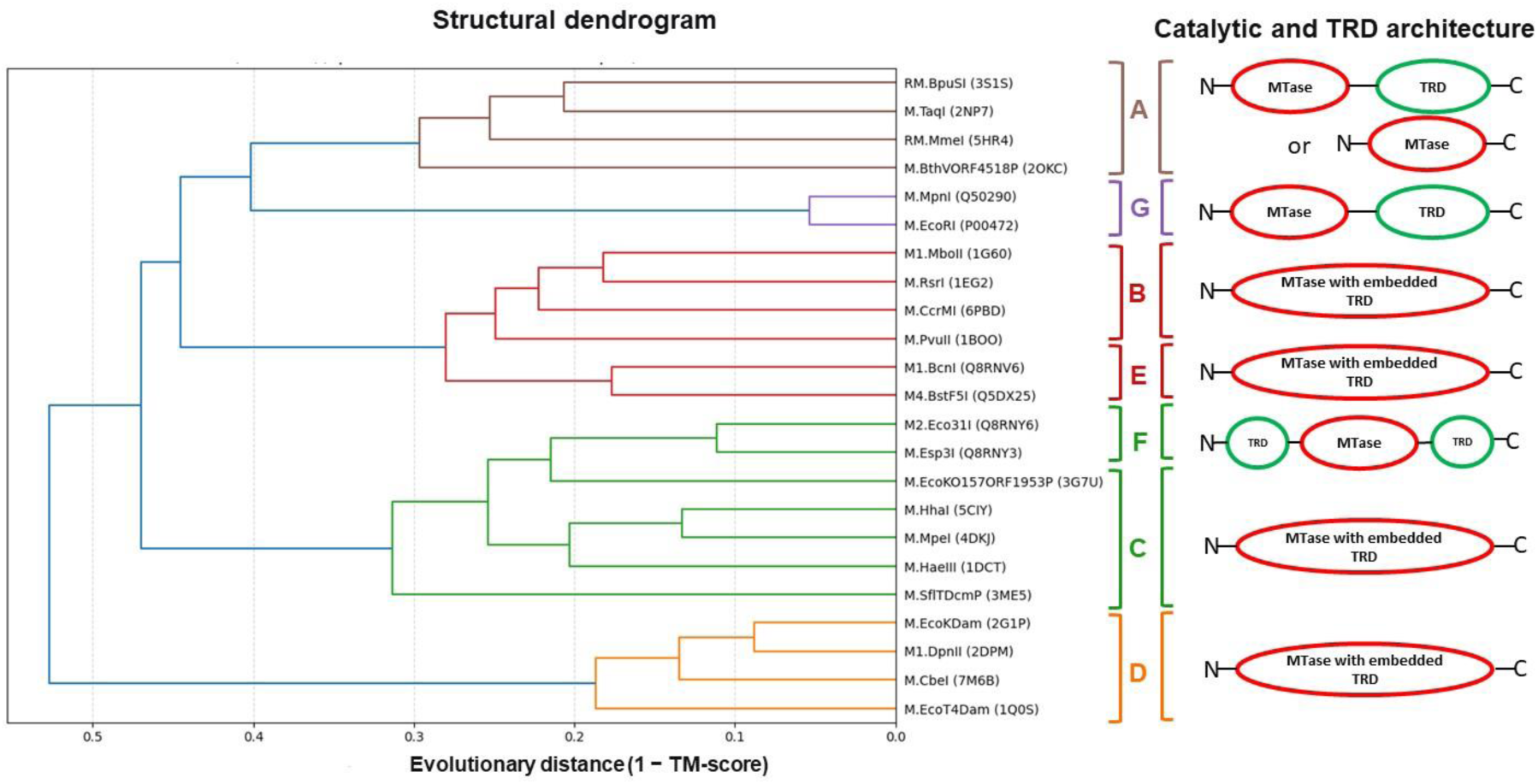
Structural relationships and domain architecture across the seven MTase topological classes (A-G). (Left) UPGMA hierarchical clustering dendrogram of representative MTase structures based on pairwise evolutionary distance (1 - TM-score), calculated from circular-permutation-aware structural alignments (jCE-CP / US-align). The vertical dashed line marks the distance threshold of 0.38, which separates the topologies into three major structural superfamilies: (i) A + G, (ii) B + E, and (iii) C + F, with topology D forming a separate branch. (Right) Schematic linear arrangement of the catalytic core (pink ovals, MTase) and target recognition domains (green ovals, TRD) along the polypeptide chain (N to C) for each topological class.

Topologies B and E are structurally related. Structural superposition of M.CcrMI (6PBD, topology B) and M1.BcnI (Q8RNV6, topology E) gave a TM-score of 0.70 and RMSD of 3.01 Å over 238 aligned residues (Figure 5a). Despite their different topologies — B has a permuted strand order, whereas E retains the canonical S0→S5→S6→S7→S4→S3→S1→S2 arrangement — the two structures superimpose well, with the alignment extending over the TRD-containing loops in addition to the catalytic core. The S0 strand at the N-terminus of M1.BcnI spatially corresponds to the S0 of M.CcrMI, consistent with a circular permutation.

**Figure 5.**
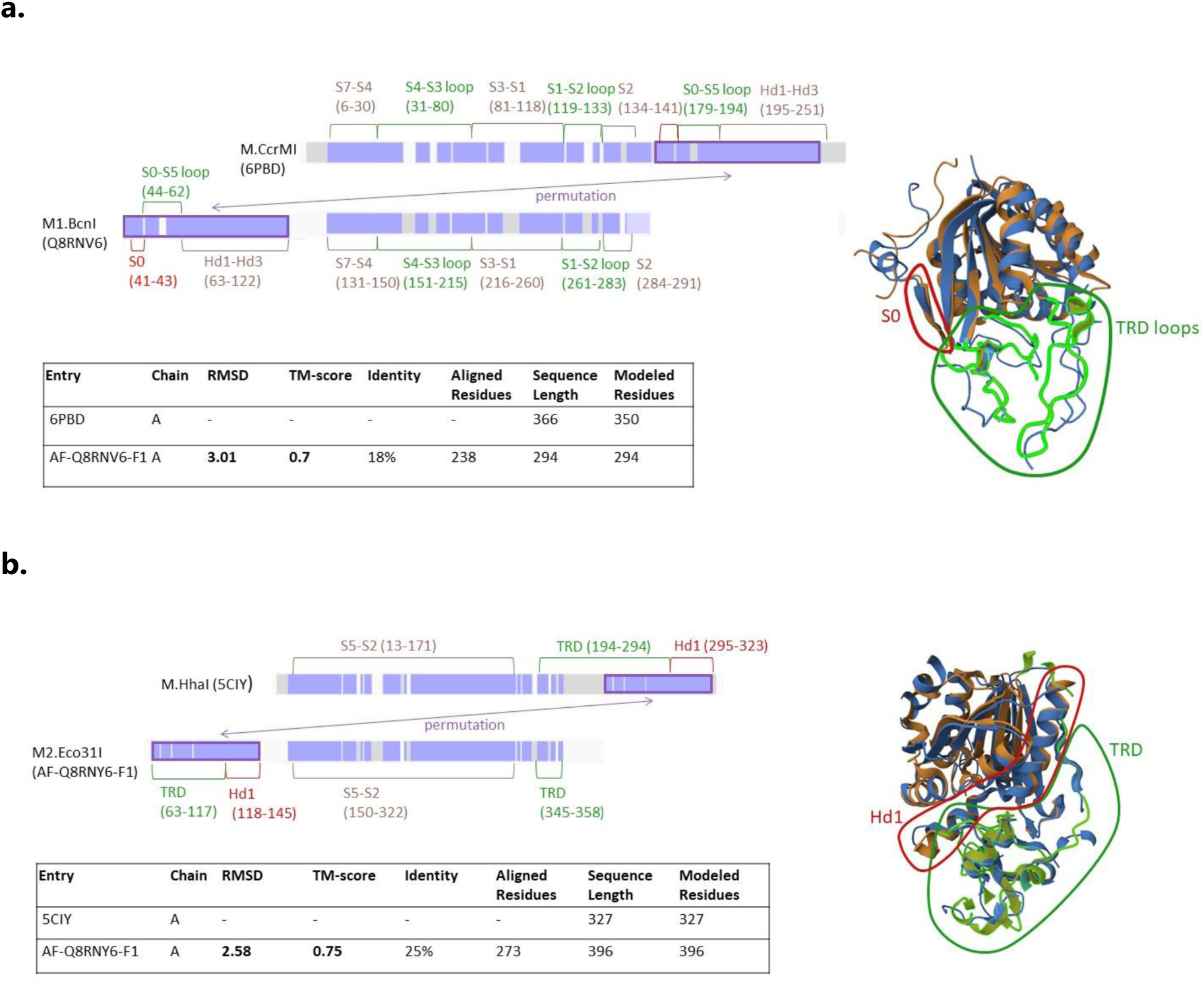
Structural relationships between topologies. (a) Comparison of M.CcrMI (6PBD, topology B) and M1.BcnI (Q8RNV6, topology E). Left: linear arrangement of structural blocks. Right: structural superposition (jCE-CP; TM-score 0.70, RMSD 3.01 Å over 238 aligned residues). The S0 strand (red) and the aligned TRD regions (green) are highlighted. (b) Comparison of M.HhaI (5CIY, topology C) and M2.Eco31I (Q8RNY6, topology F). Left: linear arrangement of structural blocks. Right: structural superposition (jCE-CP; TM-score 0.75, RMSD 2.58 Å over 273 aligned residues). The aligned TRD regions (green) and the relocated Hd1 helix (red) are highlighted. The rearrangement captured the C-terminal part of the TRD together with the Hd1 helix, while the linker and the remaining portion of the TRD stay at the C-terminus.

Topologies C and F share a homologous TRD at different positions. Structural superposition of M.HhaI (5CIY, topology C) and M2.Eco31I (Q8RNY6, topology F) gave a TM-score of 0.75 and RMSD of 2.58 Å over 273 aligned residues (Figure 5b). Both structures share the same core fold, but the rearrangement extends beyond the catalytic core: the permutation has captured the C-terminal part of the TRD together with the Hd1 helix, while the linker and the remaining portion of the TRD stay at the C-terminus. As a result, in M2.Eco31I the TRD is split into two segments — an N-terminal part (residues 63–117) and a C-terminal part (residues 345–358). The Hd1 helix is correspondingly relocated.Taken together, these results indicate that the seven topologies fall into three structural superfamilies — A + G, B + E, and C + F — with D as an outlier. Within each superfamily, the catalytic core is conserved, while peripheral elements (S0, TRD, Hd1) have undergone permutation or relocation.

### Web application for MTase topology analysis

To make the structure-based classification accessible to the research community, we developed an open-source pipeline and an interactive web application, MTase Topology Analyser (source code and Colab notebook available at https://github.com/MVolobueva/MTase_topology_analyser). The tool accepts a PDB ID, an AlphaFold (UniProt) ID, or a user-uploaded PDB file as input. It automatically scans the structure for catalytic motif IV using default patterns (DPPY, NPPY, PC, PS). Users can also specify custom motifs or, if the catalytic motif is unknown, enter a broad regular expression (e.g., [A-z]) to scan all beta-strands without motif restrictions. For each detected motif, the application reports the motif sequence and position, the number of beta-strands in the catalytic core, the linear strand order, and the residue boundaries of the core. The output includes a linear topology string, a 2D topology diagram, an interactive 3D structure with the catalytic core highlighted, and a downloadable DSSP secondary-structure assignment file. Note that the web tool performs structural core extraction and strand-order determination rather than direct assignment to topological classes A-G; final class assignment is determined from the strand order, the presence of auxiliary strands (S0 and S-1), the S6-S7 gap length, and the arrangement of terminal helices. Structures with incomplete catalytic cores (fewer than six beta-strands) or lacking recognizable motifs are reported as unassigned. The web interface is available at https://mtasetopologyanalyser.streamlit.app/.

## DISCUSSION

We propose a classification system that groups prokaryotic DNA methyltransferases into seven structural classes (A–G) based on the architecture of their catalytic domain. Within each topology, members share (i) a conserved strand order of the catalytic core, (ii) a characteristic catalytic motif, and (iii) consistent clustering in the TM-score dendrogram (Figure 4). Between topologies, these features differ, and structural superposition yields lower TM-scores. Together, these observations are consistent with a common origin within each class, suggesting that they represent families in the evolution of DNA MTases.

Structural relationships between topologies. Clustering based on TM-scores revealed three structural superfamilies: (i) A + G, (ii) B + E, and (iii) C + F, with topology D forming a separate branch (Figure 4). Within each superfamily, the catalytic core is conserved, while peripheral elements have undergone permutation or relocation.

Topologies B and E are structurally related. Structural superposition of M.CcrMI (6PBD, topology B) and M1.BcnI (Q8RNV6, topology E) gave a TM-score of 0.70 and RMSD of 3.01 Å over 238 aligned residues (Figure 5a), with the alignment extending over the TRD-containing loops in addition to the catalytic core. The S0 strand at the N-terminus of M1.BcnI spatially corresponds to the S0 of M.CcrMI, consistent with a circular permutation. Notably, M1.BcnI is only 294 residues long, with the first ∼40 residues forming an extended N-terminal loop; there is no space for a full TRD domain. Nevertheless, three recognition loops — S1–S2, S0– S5, and S4–S3 — of M1.BcnI structurally align with the corresponding loops of M.CcrMI, suggesting that target recognition in E is mediated by the same set of surface loops as in B. Thus, topology E likely represents a permuted variant of topology B in which the dedicated TRD is absent and target recognition is mediated by surface loops.

Topologies C and F share a homologous TRD, but its arrangement differs substantially. Structural superposition of M.HhaI (5CIY, topology C) and M2.Eco31I (Q8RNY6, topology F) gave a TM-score of 0.75 and RMSD of 2.58 Å over 273 aligned residues (Figure 5b), indicating that the catalytic cores of the two enzymes are highly similar. The rearrangement, however, extends beyond the catalytic core: the permutation has captured the C-terminal part of the TRD together with the Hd1 helix, while the linker and the remaining portion of the TRD stay at the C-terminus. As a result, in M2.Eco31I the TRD is split into two segments — an N-terminal part (residues 63–117) and a C-terminal part (residues 345–358). These observations are consistent with a complex domain rearrangement during evolution. Thus, topology F likely represents a structural variant of topology C in which the TRD is split into two segments.

### TRD as a mobile and heterogeneous module

Across topologies, the TRD occupies different positions and adopts different structural forms. In topology A, target recognition is mediated by a separate S subunit of the complex, which may be C-terminally fused to the MTase. In topology B, the TRD can be either an inserted domain or surface loops (e.g., M.CcrMI uses loops for recognition). In topology C, the TRD forms a single globular domain at the C-terminus. In topology D, the TRD consists of α-helices located between Hd3 and S7. In topology E, a dedicated TRD is absent, and recognition is mediated by surface loops. In topology F, the TRD is split into two segments, one N-terminal and one C-terminal. In topology G, the TRD is a separate C-terminal module.

This variability contrasts with the conserved catalytic core and suggests that TRD relocation and rearrangement have accompanied the diversification of DNA MTases. Importantly, TRDs are not necessarily homologous within a single topology — as seen in topology A, where the S subunit and fused TRDs may have different origins — and are certainly not homologous between topologies. It is therefore likely that TRDs evolved independently in different MTase lineages, rather than deriving from a single ancestral domain.

### Topology G as a distinct structural arrangement

As described above, topology G represents a distinct structural arrangement. The predicted arrangement — an antiparallel S7, an additional S8 strand, and the replacement of the canonical Hd3 helix by a β-strand — is supported by several independent features of the models, including the length of the S6– S7 segment, the presence of a proline within it, and a hydrophobic core that stabilizes the alternative fold. Nevertheless, because these observations are based solely on AlphaFold predictions, the arrangement should be treated as provisional until an experimental structure becomes available.

### Large insertions within the catalytic domain

Large insertions (>20 aa) not corresponding to known TRDs were identified in several topologies (examples in Table 2). These form surface-exposed loops that do not disrupt the αβα-sandwich core. Some inserts, particularly in topology A, exceed 100 aa. These loops may contribute to substrate specificity or regulation, but their roles remain to be determined.

**Table 2.**
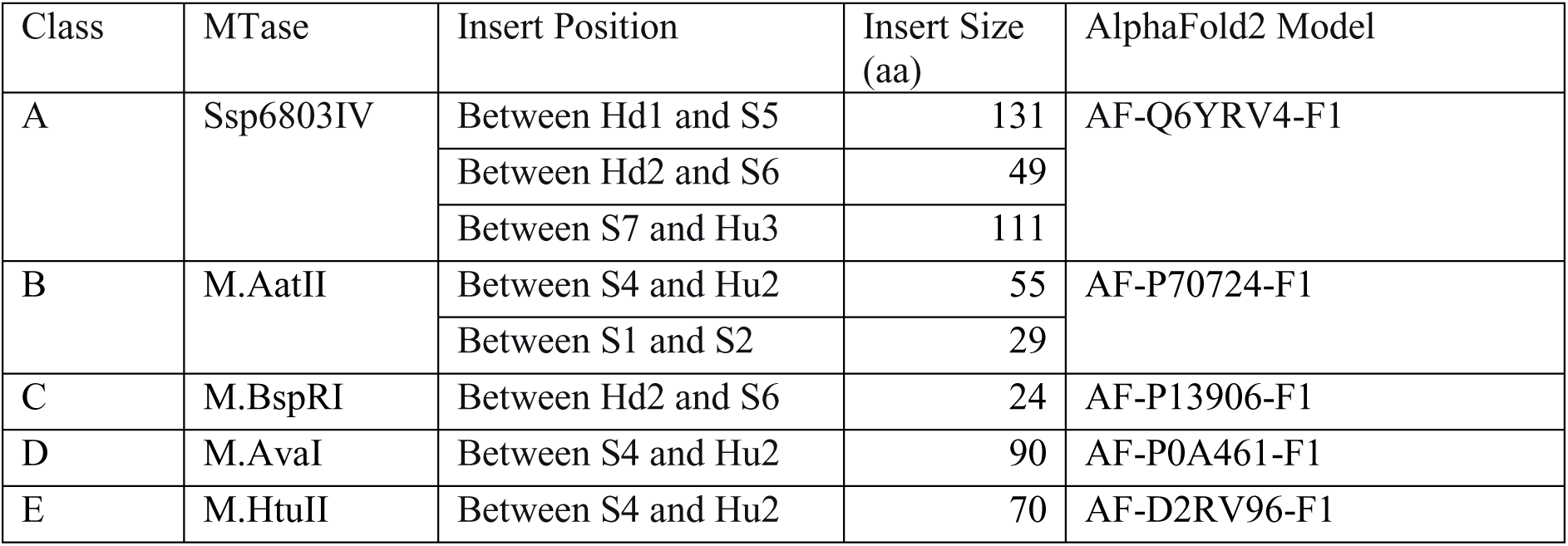
Examples of large insertions within the catalytic domain. For each example, the topological class, the MTase name, the position of the insert relative to secondary-structure elements, its size in amino acids, and the corresponding AlphaFold model are given. Insertions were defined as segments longer than 20 aa that do not correspond to known target recognition domains (TRDs).

| Class | MTase | Insert Position | Insert Size (aa) | AlphaFold2 Model |
|---|---|---|---|---|
| A | Ssp6803IV | Between Hd1 and S5 | 131 | AF-Q6YRV4-F1 |
|  |  | Between Hd2 and S6 | 49 |  |
|  |  | Between S7 and Hu3 | 111 |  |
| B | M.AatII | Between S4 and Hu2 | 55 | AF-P70724-F1 |
|  |  | Between S1 and S2 | 29 |  |
| C | M.BspRI | Between Hd2 and S6 | 24 | AF-P13906-F1 |
| D | M.AvaI | Between S4 and Hu2 | 90 | AF-P0A461-F1 |
| E | M.HtuII | Between S4 and Hu2 | 70 | AF-D2RV96-F1 |

### Comparison with previous classifications

Our system recapitulates the known groups (α, β, γ, C5) and reveals additional diversity (Table 3). Topology B corresponds to the β subtype, D to the α subtype, and C to the C5 subtype. Topology G corresponds to proteins previously assigned to the γ subtype. Topology A corresponds to γ for members with a C-terminally fused TRD, but also includes a distinct group with multi-subunit architecture — particularly Type I R-M systems — that lacks a fused TRD and relies on a separate S subunit; this architecture has no direct counterpart in the motif-based classification. Topologies E and F represent arrangements not accounted for by previous systems: E lacks a large inserted TRD and uses surface loops for recognition, while F is distinguished by its split TRD and sharing m5C activity and the PC motif with topology C.

**Table 3.** Comparison of the topological classification with the previous motif-based classification. For each new class, the methylation type, the corresponding subtype in the motif-based scheme (where applicable), and notes on the target recognition domain (TRD) are given.

| **New class** | **Methylation Type** | **Corresponding Subtype (Motif-based)** | **Notes on TRD** |
|---|---|---|---|
| **A** | m4C, m6A | γ (partial) | Correspondence to γ is valid only for members with a C-terminally fused TRD. |
| **B** | m4C, m6A | β |  |
| **D** | m4C, m6A | α |  |
| **E** | m6A |  | Lacks a large, inserted TRD; recognition likely via loops. |
| **C** | m5C | C5 |  |
| **F** | m5C |  |  |
| **G** | m6A | γ |  |

## Limitations

Four limitations should be noted.

First, the parser searches for two default motif families, [SND]P[PL][YFW] and P[CS], although the search pattern can be adjusted by the user. Two limitations follow from this design. On the one hand, the detection of the beta-sheet depends on the geometry of the region around the motif, and in some structures this geometry does not allow the sheet to be assembled correctly even when the motif itself is present. On the other hand, automated detection of MTases with motifs that deviate from the two default families requires either providing the specific motif in advance or using a broad wildcard regular expression (such as [A-z]) to scan all beta-strands. In the latter case, the pipeline outputs all candidate beta-sheets in the structure, requiring manual selection of the true catalytic core. This limitation is illustrated by three reviewed Swiss-Prot MTases (P00473, P43871, Q57168) carrying non-canonical motif IV variants (DPQY, DIPY, NPHF) and five others (P25267, P29749, P44431, P50190, Q58617) carrying canonical motifs but failing the geometric criterion (Table S3).

Second, in several experimental structures, individual β-strands were not resolved owing to crystallographic disorder (e.g., 7BTO, 7BTR, 7LN1, 9EXH), leading to incomplete catalytic domains and low-confidence or unassigned classification, although we prioritized structures with the most complete catalytic cores. This limitation applies primarily to experimental structures; for AlphaFold-predicted models, strand coverage is generally more complete, although DSSP-based secondary-structure assignment can still fail in some cases.

Third, 16 AlphaFold-predicted models were assigned to topology A manually, based on motif similarity and the absence of features characteristic of other classes, rather than by the automated pipeline. While their catalytic motifs are compatible with class A, we cannot exclude that some of these models represent previously unrecognized topologies, or alternatively that they reflect inaccuracies in the predicted structures. Resolving these possibilities would require experimental 3D structures.

Fourth, the automated classification may produce incorrect assignments in structures containing additional strands (S0, S-1, S-2, S-3), in which individual strands are unresolved, or in which the assignment of the Hd1 helix is ambiguous. In such cases, the result was verified manually by checking whether the detected catalytic motif is characteristic of the assigned class. This is particularly relevant for classes A and C, which share the same β-sheet architecture but carry entirely different catalytic motifs; when the motif is specified as input, the two classes can be reliably distinguished.

## CONCLUSION

Our classification system provides a framework for understanding the diversity and evolution of prokaryotic DNA MTases. Its applications are likely to benefit genomic annotation, evolutionary studies, and the engineering of novel MTases for biotechnological applications. The present classification is restricted to prokaryotic enzymes; extending it to eukaryotic DNA MTases will be addressed separately.

## DATA AVAILABILITY

All experimentally determined structures analyzed in this study are available from the PDB un der the accession codes listed in Table S1a. AlphaFold-predicted models are available from t he AlphaFold Database (https://alphafold.ebi.ac.uk/). The structure-based classification pipeli ne (DSSP, strand-order analysis, topology assignment) is available at:

- Source code: https://github.com/MVolobueva/MTase_topology_analyser
- Web interface: https://mtasetopologyanalyser.streamlit.app/
- Colab notebook: https://colab.research.google.com/github/MVolobueva/MTase_top ology_analyser/blob/main/MTase_Topology_Analyzer.ipynb

All source code is released under an open-source licence (MIT).

## SUPPLEMENTARY DATA

Supplementary Data are available https://drive.google.com/drive/folders/17yOGEkqMKVI1TD7ovUX67N2pSsPxDsZv?usp=sharing

## AUTHOR CONTRIBUTIONS

M. Samokhina conceived the study, performed the formal analysis, developed the classification system, implemented the algorithm, conducted the investigation, and wrote the original draft. A. Alexeevski supervised the research, provided conceptual guidance, acquired funding, and reviewed and edited the manuscript.

## ACKNOWLEDGEMENTS

The authors thank Ivan Rusinov for his valuable recommendations on the manuscript content and his assistance with text refinement.

